# Microparticle-delivered Cxcl9 prolongs Braf inhibitor efficacy in melanoma

**DOI:** 10.1101/2022.05.24.493271

**Authors:** Gabriele Romano, Francesca Paradiso, Peng Li, Pooja Shukla, Lindsay N Barger, Olivia El Naggar, John P Miller, Roger J Liang, Timothy L Helms, Alexander J Lazar, Jennifer A Wargo, Francesca Taraballi, James C Costello, Lawrence N Kwong

## Abstract

Patients with BRAF-mutant melanoma show significant responses to combined BRAF and MEK inhibition, but most relapse within 2 years. A major reservoir for drug resistance is minimal residual disease (MRD), comprised of drug-tolerant tumor cells laying in a dormant state. Towards exploiting potential therapeutic vulnerabilities of MRD, we established a genetically engineered mouse model of Braf^V600E^-driven melanoma MRD wherein genetic Braf^V600E^ extinction leads to strong but incomplete tumor regression. Transcriptional time-course analysis after Braf^V600E^ extinction revealed that after an initial surge of immune activation, tumors later became immunologically “cold” after MRD establishment. Computational analysis identified candidate T-cell recruiting chemokines that may be central players in the process, being strongly upregulated initially and steeply decreasing as the immune response faded. Therefore, we hypothesized that sustaining the chemokine signaling could impair MRD maintenance through increased recruitment of effector T-cells. We show that intratumoral administration of recombinant Cxcl9, either naked or loaded in microparticles, significantly impaired MRD relapse in BRAF-inhibited tumors, including several complete responses after microparticle-delivered rCxcl9 combined with BRAF and MEK-inhibition. Our experiments constitute a proof of concept that chemokine-based microparticle delivery systems are a potential strategy to forestall tumor relapse and thus improve the clinical success of frontline treatment methods.

## INTRODUCTION

BRAF-mutant melanoma has become an archetype of targeted therapy after the successful advent of BRAF and MEK inhibitors (BRAFi and MEKi) in clinical practice. Melanoma patients carrying the BRAF^V600E^ alteration, in particular, show impressive responses to combinations of BRAFi+MEKi, a therapeutic strategy that extends the lives of thousands of patients every year(1). Nevertheless, most patients with melanoma show signs of relapse within the first 2 years after treatment start(1). The most frequent molecular mechanism of drug resistance is MAPK pathway reactivation through events such as BRAF amplification or the acquisition of additional MAPK-activating alterations(2). The occurrence of drug resistance and the eventual relapse can be explained by the survival of a cadre of tumor cells that eventually leads to tumor re-growth. These drug-tolerant cells constitute the Minimal Residual Disease (MRD), where cancer cells linger in a dormant state(3).

MRD can be fueled by intrinsic features of cancer cells or extrinsic mechanisms of the microenvironment. Among the extrinsic factors, one of the most relevant is the immune system. When targeted therapy is administered, a strong immune activation is triggered(4). BRAFi have been demonstrated to induce the expression of melanoma antigens, favoring the recognition by the immune system and the infiltration of CD8 effector and CD4 helper T-cells(5). In this regard, the magnitude of the mounted response is key to disease eradication. If a few cells manage to escape the initial immune response, they can fuel the establishment of MRD. Consistently, the patients with the most favorable long-term responses to BRAFi+MEKi are associated with a baseline elevated immune infiltrate, a status described as “hot” tumor bed(6–9). Contrariwise, the “cold” immune microenvironment is among the adverse prognostic factors of targeted therapy. A cold environment has a high abundance of suppressive cells and/or a low percent of effector and helper T-cells(10). In such a microenvironment, MRD has an increased possibility of surviving during treatment and, over time, acquiring the features to relapse to a full-blown tumor. This is true also for other therapeutic approaches, including immune checkpoint inhibitor therapy(10).

Converting a “cold” tumor microenvironment to a “hot” one is a potential mechanism to impair the establishment of MRD and prolong the clinical success of targeted therapy. Recruiting more effector cells to the tumor site can be a viable strategy for patients with low baseline immune infiltrate. In this work, we present a microparticle-based approach to favor the recruitment of Cd8+ T cells into melanoma MRD using the computationally identified Cxcl9 chemokine. We demonstrate that the administration of Cxcl9 induces an ingress of Cd8+ T cells into the tumors and that, if administered concomitantly with BRAFi, it significantly delays the occurrence of tumor relapse.

## RESULTS

### A mouse model of melanoma Minimal Residual Disease

As a Minimal Residual Disease (MRD) model, we utilized our previously described inducible and conditional GEMM model of Braf^V600E^ mutant melanoma, iBIP(9). Briefly, topical 4-hydroxy-Tamoxifen (4-OHT) and systemic doxycycline (dox) administration restrict Braf^V600E^ expression and Pten and Cdkn2a knockout to melanocytes. These develop into fully formed melanomas in 6-8 weeks. Dox withdrawal causes Braf^V600E^ extinction (hereafter “Braf extinction”) and significant tumor regression, consistent with the driving oncogenic role of Braf^V600E^ (**Fig. 1A**). In nearly all cases (>90%), the regressed tumors do not entirely disappear but stall in a state we describe as MRD (**Fig. 1A-B**), as the tumors became impalpable. Moreover, we verified that the tissue contains residual tumor cells through IHC staining of GFP, a built-in tumor marker (**Fig. S1**). Notably, the MRD is established around 30 days after Braf extinction and can be re-triggered with dox-induced Braf re-expression to cause a rapid tumor relapse, even after prolonged extinction (**Fig. 1A**). Overall, this model mimics patients with Braf^V600E^ melanoma who initially respond to BRAFi, but who eventually relapse due to BRAF overexpression, BRAF amplification, or the expression of a non-targetable BRAF splice isoform (collectively, >30% of human patient BRAFi resistance mechanisms) (2,11).

**Figure 1.**
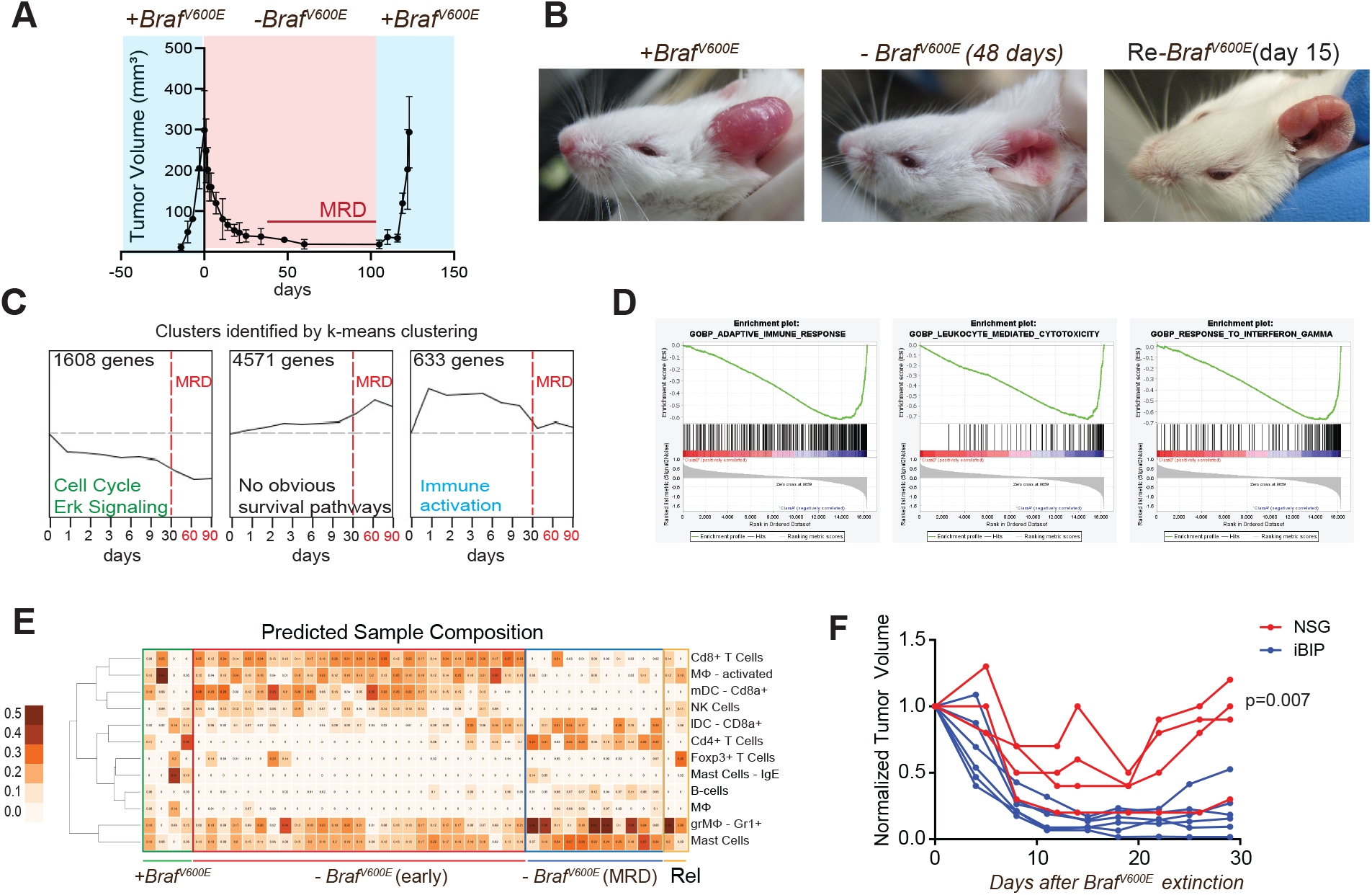
A GEMM model of melanoma reveals that Minimal Residual Disease (MRD) is a “cold” immune microenvironment. A) Tumor growth curve of iBIP mice treated with Doxycycline (dox) administration, after Braf extinction, and after Braf re-induction. B) Representative pictures of tumor-bearing iBIP mice on dox, after Braf extinction, and after Braf re-induction. C) Top 3 enriched pathways in early Braf extinction vs. MRD according to Gene Set Enrichment Analysis (GSEA) analysis. D) k-means clustering of microarray data of 45 tumors. Three clusters of interest are reported, and the number of genes in the cluster is specified. Indicated days are after BRAF extinction (dox withdrawal). E) Immune deconvolution analysis (GEDIT) predicting the immune infiltrate composition in iBIP tumors on dox, during early BRAFi (<30d), during MRD (>30d), and in Braf re-expression-induced relapsing tumors (Rel). Boxes on the x-axis represent single samples. Heatmap values represent predicted populations’ abundance. F) Tumor growth curves of Braf-extinguished iBIP tumor xenografts in immunocompetent mice (iBIP) or immune-deficient mice (NSG). The indicated p-value is calculated in an unpaired t-test between groups 30d after tumor extinction.

### Transcriptomic time-course analysis of MRD-establishment

To investigate the molecular mechanisms underlying the establishment of MRD, we performed a time-course transcriptional analysis on iBIP tumors after Braf extinction. We collected 45 tumors from as early as 8h to as late as 160d after dox withdrawal.

Using RNA microarrays, we analyzed differentially regulated genes before and after MRD establishment (30d after Braf extinction) using the Short Time-series Expression Miner (STEM)(12) algorithm and detected multiple trends of interest (**Fig. 1C** and **Table S1**). The top-scoring gene sets downregulated in MRD, as expected, were ERK and proliferation pathways, reflecting the facts that MRD is quiescent and that Braf extinction impairs the MAPK pathway. Notably, we did not identify pathways significantly upregulated in MRD that would obviously contribute to cell survival or to immune suppression/exhaustion (e.g., PD1, PD-L1, CTLA-4); instead, the top pathways related to neural and skeletomuscular genes, possibly reflecting the larger relative contribution of normal neurons and muscle cells in the small MRD mass. Most intriguingly, we observed a strong enrichment for highly upregulated immune signature gene sets as early as 8h after Braf extinction, until 9 days when it then steeply declined to baseline upon MRD establishment (at ~30d, **Fig. 1C**). To dissect the involved immune pathways more finely, we performed GSEA analysis, which showed that the top 25 enriched pathways in the early phase of Braf extinction (before ~30d) were related to antigen presentation, lymphocyte homing, and activation (**Fig. 1D**). Consistently, immune cells deconvolution analysis (GEDIT) predicted an early influx of Cd3+ and Cd8+ T Cells and myeloid antigen-presenting cells, which faded away when MRD was established (**Fig. 1E**).

To determine whether the immune response to Braf extinction is critical to the anti-tumoral effect, we inoculated iBIP-derived tumor cells into either iBIP (immunocompetent) or NSG (severely immunodeficient) mice. When doxycycline was withdrawn, NSG mice showed impaired tumor shrinkage and increased spontaneous relapse, consistent with the hypothesis that MRD relapse is at least in part regulated by the immune microenvironment (**Fig. 1F**).

### Cxcl9 is upregulated upon Braf extinction/inhibition and mirrors the overall immune infiltrate dynamics

To nominate central regulators of the immune response to BRAFi, we adapted our published TRAP network analysis algorithm(9), designed to identify genes, in this case cytokines, that are most central to a set of biological processes by querying an extensive compendium of mouse expression datasets. Chemokines are a large category of molecules involved in the chemoattraction, activation, and differentiation of immune cells; thus, we hypothesized that they are central regulators of the immune response to BRAFi. Using the k-means clustering result (**Fig. 1D**), we identified 68 significantly enriched gene sets in the “immune activation” cluster (**Table S2**). We queried these 68 gene sets in the TRAP network to establish a subnetwork of cytokine-to-immune pathway relationships (**Fig. 2A**). The output of the TRAP analysis on the subnetwork is a weighted degree centrality score for each cytokine. We reasoned that cytokines that are initially highly upregulated upon BRAF extinction, then highly downregulated in MRD, and with a high degree centrality score, are high-confidence candidate immune regulators. We, therefore, plotted these elements together: high centrality score (**Fig. 2B**, circle size), significant downregulation from early BRAF extinction to MRD (**Fig.2 B**, negative log2 fold-change and high statistical significance), and high upregulation from control to early BRAF extinction (**Fig. 2B**, circle color) (**Table S3)**. Two of the top candidates, Cxcl9 and Cxcl10, are highly related paralogs whose protein products bind Cxcr3, are IFN-gamma response genes, and are well-described as important for Cxcr3+ T cell attraction and/or activation, including in melanoma (13). As shown in **Fig. 2C**, their expression pattern closely mirrored the overall immune score trend. Notably, upon BRAF re-expression-induced tumor relapse, the immune score and the Cxcr3 ligands remained low (**Fig. 2C**, “Relapse”). We note that a third paralog of the same family, Cxcl11, showed similar parameters and trends to Cxcl9 and Cxcl10, except for a lower network centrality.

**Figure 2.**
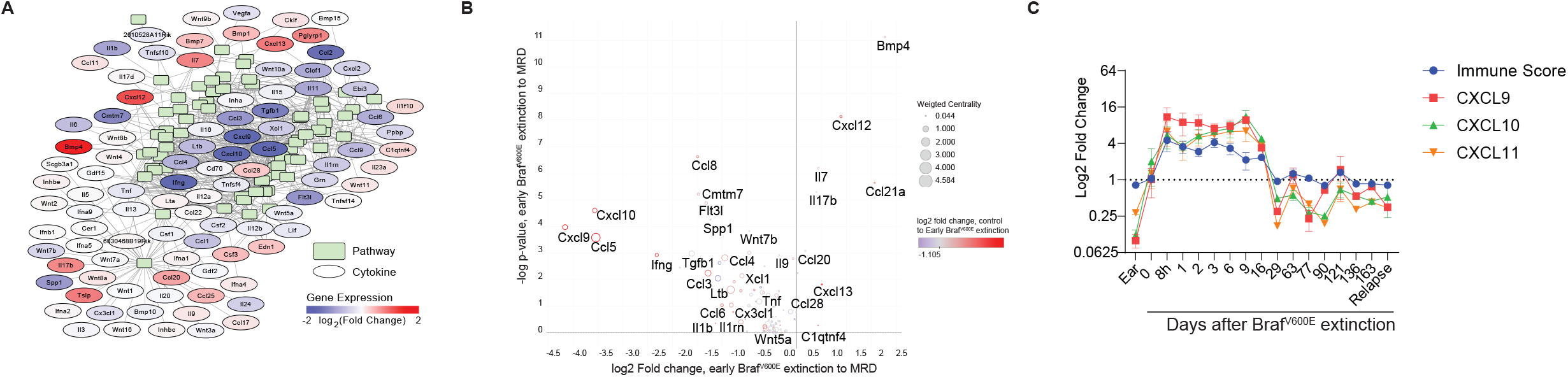
Cxcr3 ligands are key players in the immune response to BRAF extinction. A). Visual representation of cytokine-to-pathway connections of the TRAP network. Cytokines (ovals) and relevant associated pathways (rectangles) are shown. Color coding for chemokines represents log2 fold change in MRD vs. Early Braf extinction. B) Volcano plot representing the log2 fold change from early Braf extinction to MRD of the indicated chemokines (x-axis), and the corresponding log2 p-value (y-axis). The size of the circles represents centrality calculated by TRAP, and the color of the circle represents the log2 fold change from control to early Braf extinction. Only the 102 highest-centrality genes are shown for clarity. C) Cxcl9, Cxcl10, and Cxcl11 expression over time compared with the overall immune signature.

We then selected Cxcl9 as a representative Cxcr3 ligand for initial in vivo validation, as it had the largest and most significant fold-change of the 3 top candidates (**Fig. 2B**). We first validated Cxcl9 protein expression using IHC, confirming an increase upon Braf extinction and a subsequent decrease over time (**Fig. 3A-B**). We also confirmed concurrent Cd8+ and Cd4+ T-cell infiltration and abatement, with Cd8+ T cell dynamics lagging slightly after Cxcl9, consistent with a potential regulatory association (**Fig. 3B**).

**Figure 3.**
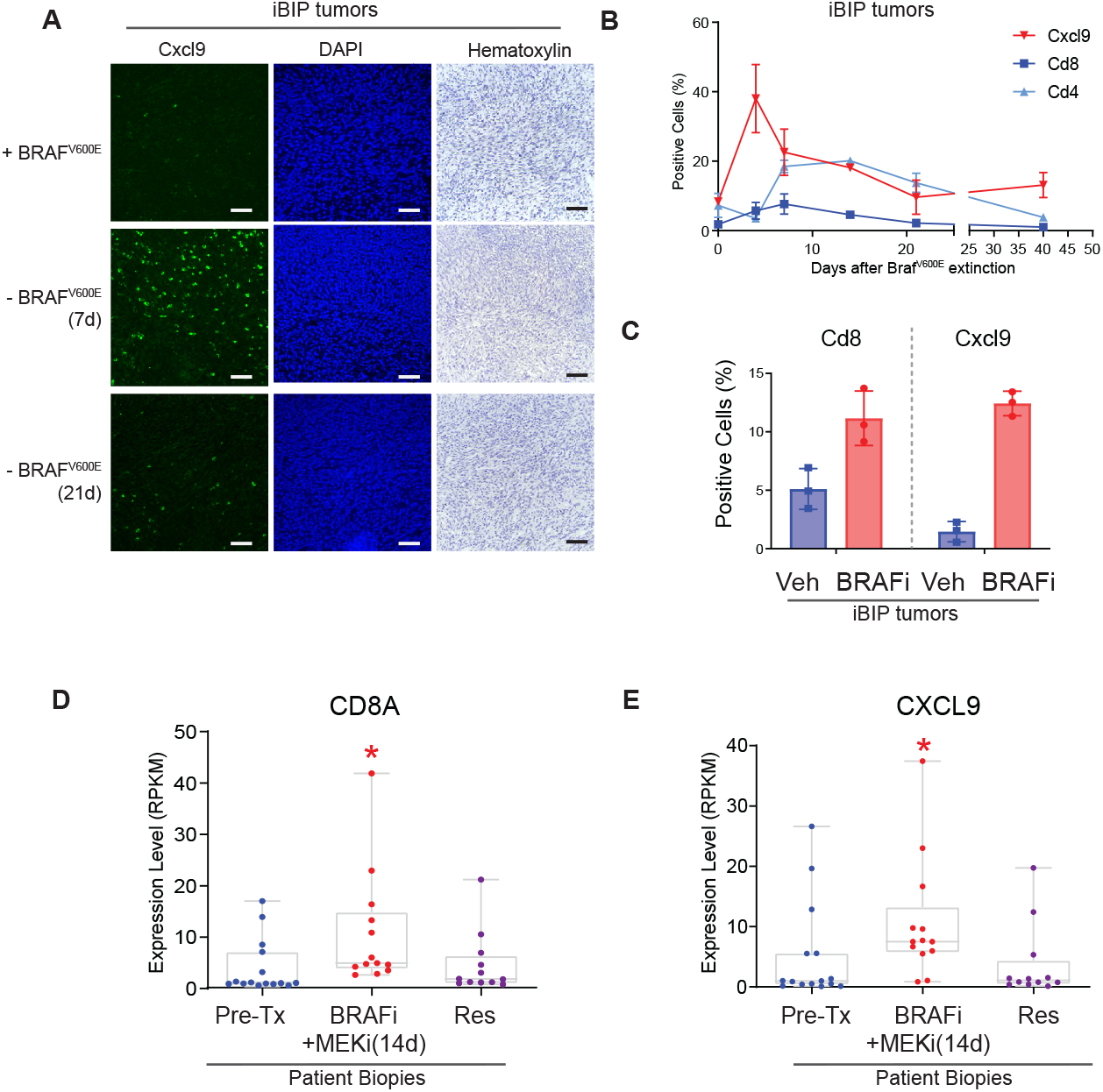
CXCL9 is induced by genetic and pharmacologic BRAF inhibition in mouse and human tumors. A) Representative immunofluorescences (IFs) for Cxcl9 protein in iBIP sections, before and after dox withdrawal. DAPI nuclear staining and Hematoxylin staining are shown. Bars represent 100 μm. B) IF time-course quantification of Cxcl9, Cd4, and Cd8 positive cells in iBIP sections after dox withdrawal (n=5, per time point). The y-axis is the % of positive cells over total cells counted in the section. C) IF quantification of Cxcl9 and Cd8 positive cells in iBIP tumor sections after pharmacologic BRAFi (PLX4720, 417 parts per million, ppm, 1 week of treatment, n=5). D) CXCL9 and E) CD8 mRNA levels in melanoma patients pre-treatment (Pre-Tx), on BRAFi+MEKi, and resistant (Res) to the treatment, as measured by RNAseq(9). Each point represents a patient sample (Pre-Tx n=15, On BRAFi+MEKi n=13, Res n=12). RPKM, Reads per Kilobase Million. * represents a p-value <0.05 calculated in an unpaired t-test compared to control.

We next asked whether Cxcl9 is also induced by pharmacological BRAFi. In both the GEMM iBIP model (**Fig. 3C**) and the previously published syngeneic “BP” model(14)(**Fig. S3**), Cxcl9 and Cd8 markers increased in the tumors in response to the BRAF inhibitor PLX4720, consistent with our genetic BRAF extinction results. To confirm clinical relevance, we analyzed our previously published human patient sample RNAseq dataset(9), which includes pre-treatment, on-treatment, and BRAFi-resistant biopsies. Similar to our mouse models, both CXCL9 and CD8A expression sharply rose on BRAFi treatment and abated upon the acquisition of BRAFi resistance (**Fig. 3D-E**).

### Recombinant Cxcl9 administration chemoattracts Cd8 lymphocytes in vitro and in vivo

We next reasoned that one or all three of these Cxcr3 ligands might regulate T-cell tumor infiltration and/or activation in response to BRAF inhibition. First, we verified the ability of recombinant mouse Cxcl9 (rCxcl9) to attract Cd8+ mouse lymphocytes in vitro using a transwell assay. As shown in **Fig. 4A**, Cxcl9 significantly induced Cd8+ T cell chemoattraction in a dose-dependent manner, and similarly to its paralogs Cxcl10 and Cxcl11 **(Fig. 4A)**. Interestingly, Cxcl9 and Cxcl11, but not Cxcl10, also slightly induced activation markers in T Cd8+ cells (Cd69, early and Cd25, late activation marker, **Fig. S2**).

**Figure 4.**
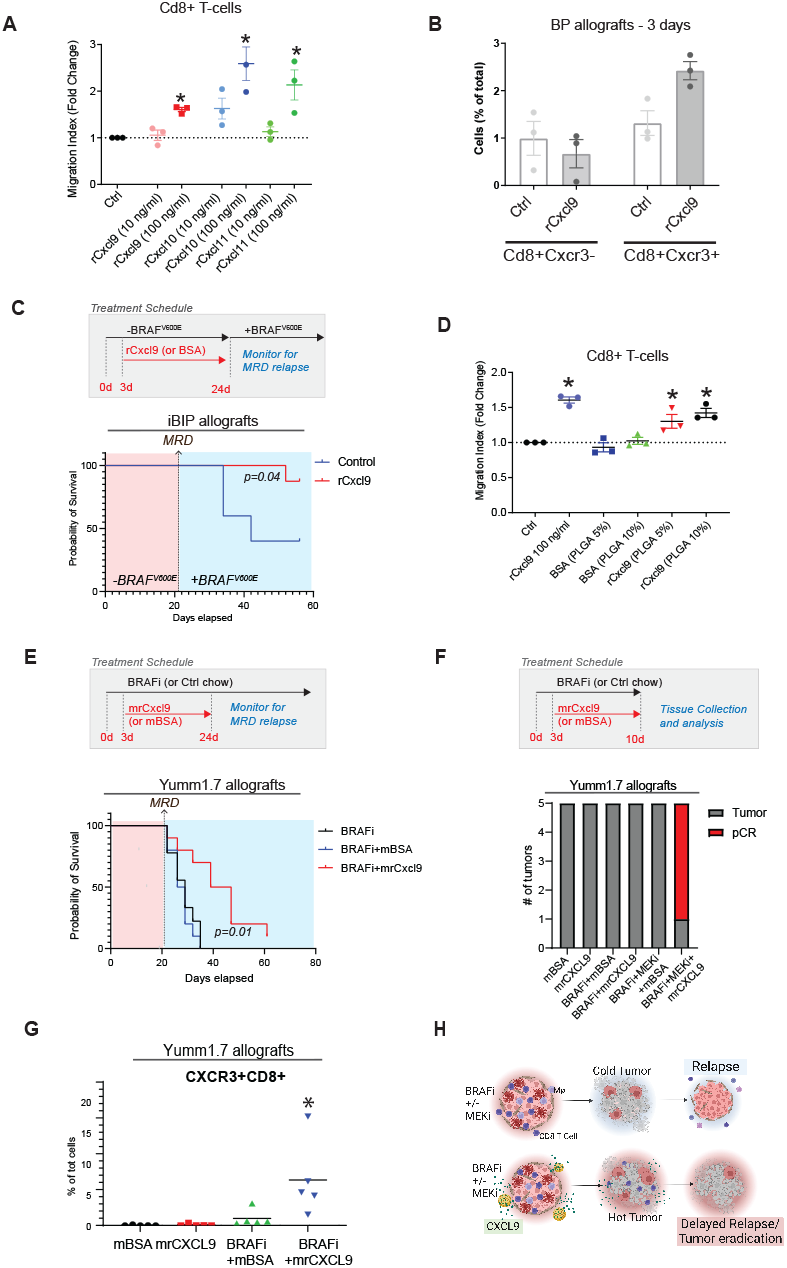
Recombinant Cxcl9 attracts Cd8 T cells in vitro and in vivo and delays the occurrence of tumor relapse. A) Transwell migration assay of mouse Cd8+ T-cells using chemokines at the indicated concentrations. The migration index is calculated as the number of cells migrated over the total number of cells. B) Flow Cytometric analysis of Cd8+Cxcr3+ and Cd8+Cxcr3-T cell abundance in “BP” tumors after intratumoral injection of rCxcl9 (10μg/administration, for 3 days; n=3) or vehicle. The percentage of Cd8+Cxcr3+ (or Cd8+Cxcr3-) cells is calculated over the total number of cells in each sample. C) Survival curves of iBIP allografts injected with rCxcl9 or vehicle (Ctrl, n=5, rCxcl9, n=7) after dox withdrawal. At MRD establishment (21d), tumors were rechallenged with dox and observed for relapse. A death event is considered as the first tumor doubling after MRD acquisition. D) Transwell migration assay for mCd8+ T cells using BSA- or rCxcl9-loaded PLGA microparticles. The migration index is calculated as the number of cells migrated over the total number of cells. E) Survival curves of Yumm1.7 tumors treated with PLX4720 with or without microparticles that contained BSA or rCxcl9 (n=10). A death event is considered as the first tumor doubling after MRD acquisition. The vehicle control group with no BRAFi needed to be euthanized before BRAFi and therefore cannot be represented on this graph. F) Complete pathologic responses (pCR) vs live tumor tissue detected by a certified pathologist in the indicated groups. G) Immunofluorescence quantification of Cd8+Cxcr3+ T cell abundance in Yumm1.7 tumors treated with PLX4720 with or without microparticles that contained BSA or rCxcl9 (n=5). The percentage of Cd8+Cxcr3+ cells is calculated over the total number of cells in the tissues. H) Schematic of the rational treatment approach proposed in the present work. * represents a p-value <0.05 calculated in an unpaired t-test compared to control. The p-values indicated on the Kaplan-Meier curves are calculated through a Log-rank (Mantel-Cox) test.

We then tested Cxcl9 chemoattraction in vivo. We injected rCxcl9 (10ug/administration) intratumorally for 3 days and then analyzed the abundance of the infiltrating Cd8+ T cell population. As expected, rCxcl9 specifically recruited Cd8+Cxcr3+ cells into the tumors, consistent with Cxcr3 as the known binding receptor of Cxcl9 (**Fig. 4B**). To validate the therapeutic potential of our findings, we tested the effect of rCxcl9 administration in iBIP allografts. We injected rCxcl9 intratumorally (0.5-1μg/administration, QD) for 3 weeks after dox withdrawal. After MRD establishment, we re-administered dox and monitored tumors for re-growth. Notably, the control group relapsed significantly more and earlier than the rCxcl9-treated group over the observation period (**Fig. 4C, Fig. S4)**.

### A microparticle-based approach for chemokine delivery

A potential hurdle to translating Cxcl9 into therapy is that chemokines are rapidly degraded in vivo. Nanoparticles for drug delivery applications have been developed to overcome some of the limitations of free therapeutics, including drug stability and release kinetics(15,16). Toward this aim, we leveraged an approach in which rCxcl9 is encapsulated in microparticles composed of a silica core and a poly(DL-lactide-co-glycolide) acid (PLGA) outer shell (**Fig. S5**)(16). PLGA-silica particles assure chemokines protection from degradation while providing a steady release and supply.

In vitro experiments confirmed that PLGA microparticles release rCxcl9 efficiently and attract Cd8+ T mouse lymphocytes (**Fig. 4D**) to levels similar to that of naked rCxcl9. Next, we utilized the well-established Yumm1.7 syngeneic melanoma model(17) to further confirm our initial findings in the iBIP model. Yumm1.7 tumors have the same clinically relevant genetic alterations as iBIP (Pten^-/-;^ Cdkn2a^-/-^; Braf^V600E^), sharply respond to pharmacologic BRAFi, and establish MRD with a natural eventual tumor relapse in syngeneic immunocompetent C57BL6/J mice hosts.

To determine whether rCxcl9-loaded PLGA microparticles (hereafter “mrCxcl9”) affect melanoma relapse, we induced Yumm1.7 tumors and divided mice into 4 groups (n=10 per cohort): Control Vehicle (Ctrl), BRAFi, BRAFi + PLGA microparticles loaded with BSA (BRAFi+mBSA), and BRAFi + PLGA microparticles loaded with rCxcl9 (BRAFi+rCxcl9). Between the PLGA tested concentrations (10% or 5% PLGA 50:50), 10% was chosen for translation in vivo since it induced the highest migration of T CD-8 cells in vitro. Starting 3 days after BRAFi, the two PLGA groups were injected intratumorally with microparticles for 3 weeks, once a week, at 50ug of total rCxcl9 or BSA per administration. All groups were continuously maintained on BRAFi and monitored for tumor relapse, which we defined as the first tumor size doubling after MRD establishment. As expected, the vehicle control group with no BRAFi needed to be euthanized before 21d after tumor cell injection. By contrast, BRAFi and BRAFi+mBSA groups relapsed after a median of 29 and 27.5 days after BRAFi start, respectively (**Fig. 4E, Fig S6)**), indicating no effect of the microparticles themselves. Notably, the BRAFi+mrCxcl9 group showed a significantly delayed relapse (median 43.5 days, **Fig. 4E, Fig S6**), supporting a positive therapeutic effect of mrCxcl9 and the feasibility of microparticle delivery.

As BRAFi+MEKi is the standard of care for BRAF^V600E^ melanomas, we also tested mrCxcl9 with this inhibitor combination (PLX4720+MEK162) in the Yumm1.7 model. Strikingly, the administration of mrCxcl9 with BRAFi+MEKi induced a complete pathological response (pCR) in 4/5 tumors (**Fig. 4F, Fig. S7**) and widespread necrosis in the remaining tumor after only 10d of treatment (7d of mrCxcl9). By contrast, BRAFi+MEKi alone, or BRAFi+mrCxcl9, while potent, left detectable tumor cells in all mice, as expected. As insufficient tumor tissue remained in BRAFi+MEKi+mrCxcl9 treated mice for immunohistochemical assessment, we next took advantage of BRAFi-treated mice (without MEKi) at 10 days of treatment to assess Cd8 and Cxcr3 by multiplex immunofluorescence. Consistent with enhancing immune infiltration after BRAFi, mrCxcl9 administration on top of BRAFi caused a significant further increase of both Cd8+Cxcr3+ and overall CD8+ T-cells (**Fig. 4G**, **Fig. S8-9**).

## DISCUSSION

In this study, we demonstrated that Cxcl9 is a key immune modulator of the BRAFi-induced MRD state, which in turn manifests as a combination therapeutic modality to forestall tumor relapse (**Fig. 4H**). We leveraged the protection of PLGA-silica microparticles for rCxcl9 delivery, enabling slow intratumoral release over time, which both reduces the frequency of administration and provides a consistent output. This discovery spurs from the transcriptional and phenotypic characterization of a GEMM model of melanoma MRD. We utilized our previously published TRAP algorithm to prioritize key chemokines potentially regulating MRD establishment and maintenance, which was then validated in pharmacological, immunocompetent BRAF inhibition models.

As we did not detect an enrichment for specific immune-suppressive/evasive signals to target in the MRD, such as PD-1 or CTLA4, we hypothesized instead that sustaining T-cell recruiting signaling throughout BRAFi administration could be a viable strategy to convert MRD into immunologically “hot.” Indeed, intratumoral rCxcl9 in combination with BRAFi not only increased the influx of Cxcr3+Cd8+ T cells into the tumor but also overall Cd8+ T-cells, suggesting a positive immunologic feedback loop leading to enhanced BRAFi and BRAFi+MEKi efficacy.

While we selected Cxcl9 as our proof-of-principle chemokine, we also identified a host of other chemokines whose temporal expression profiles suggest a similar immune regulatory role; specifically, our evidence suggests their downregulation may help create the immunologically “cold” state of MRD with a low abundance of CD8+ T-cells. Of particular interest are the CXCL9 paralogs CXCL10 and CXCL11, all three of which are IFN-gamma-induced chemokines known to regulate T-cell homing through their CXCR3 cognate receptor(13). Interestingly, despite a high degree of amino acid identity, Cxcl9, Cxcl10, and Cxcl11 are known to have different roles in inflammation, graft versus host disease, and cancer(18). Consistent with this, we found differing effects of the 3 proteins on Cd8+ T cell phenotypes in vitro, with all three showing chemoattraction but Cxcl10 showing less T cell activation. Thus, we hypothesize that future testing of combinations of the 3 CXCR3 ligands and/or other high-scoring cytokines may show improved anti-tumor efficacy. Because of their versatile properties, the PLGA-silica microparticle approach we utilized is ideally suited for these future directions. Silica nanostructures can be tailored during manufacturing, changing their size, shape, porosity, and pore size to efficiently load a wide range of small and large biomolecules. Also, they can be incorporated into a wide range of synthetic polymers to finely tune the release of cytokines (19). In addition, PLGA-mp tolerability in vivo has been published extensively in different model systems, such as cancer and inflammatory diseases (19). Notably, the polymer used in this work, PLGA, is approved by the Food and Drug Administration to use in drug delivery systems, positioning PLGA-based approaches as promising candidates for future clinical trials.

Our results showing significantly prolonged treatment efficacy after microparticle-delivered rCxcl9 are consistent with the anti-tumor effect of Cxcr3 ligands seen in previous studies (20–23). However, unlike the previous efforts, which focused on rapidly growing tumors, our study recontextualizes the efficacy of Cxcr3 ligands in preventing or neutralizing drug-induced MRD, expanding their utility towards this critical clinical obstacle. In addition, this work is the first to show the chemokine therapy effect in a melanoma or MRD setting and the first to demonstrate the utility of a microparticle-based delivery.

Furthermore, the combination of the standard-of-care BRAFi+MEKi therapy with mrCxcl9 induced complete pathological responses in 4 out of 5 animals, suggesting a robust synergy. Therefore, we propose that CXCR3 ligands are clinically relevant to targeted therapies, particularly as CXCL9 expression is strongly induced in patients with melanoma treated with BRAFi but returns to baseline in patients showing relapse, mirroring CD8 expression dynamics and the observations in our mouse models. Consistent with this, multiple studies have found that CXCR3 ligands are critical for the action of anti-PD1/PDL1 immune checkpoint therapies (24,25).

Our findings demonstrate that chemokine-based approaches are a feasible strategy to regulate the immune infiltrate composition of tumors, particularly if coupled with an efficient delivery system. Immunologically “cold” tumors are, in fact, a widespread clinical problem and especially relevant to current checkpoint inhibitor therapies(4,10). Indeed, as the combination of targeted and immune approaches is experiencing the beginnings of clinical validation(26,27), there is evidence showing that baseline immune “cold” tumors can remain refractory(10,28). Chemokine-based approaches may be ideally suited to solving such issues by converting tumors to an immune “hot” microenvironment for subsequent targeted and immune checkpoint therapies to create synergy. Interestingly, several studies suggest that BRAFi+MEKi administration may negatively influence subsequent immunotherapy outcomes (29,30); further mechanistic assessment of how MRD becomes immunologically “cold” may therefore shed further light on how this might longitudinally impact the immune microenvironment. Overall, our results demonstrate that fine-tuning the composition of the immune infiltrate can be a viable adjuvant approach to boosting existing and experimental treatment approaches and eventually improving their therapeutic outcome.

