## Supplementary material for "Microparticle-delivered Cxcl9 prolongs Braf inhibitor efficacy in melanoma": Methods, Supplementary Legends

**SUPPLEMENTARY FILES**

**MATERIALS and METHODS**

*Genetically Engineered Mouse Models, mouse allografts, and tumor treatments*. The iBIP model is analogous to the previously published model(1), except for the swapping in of a conditional floxed Ink/Arf locus for the constitutive Ink/Arf knockout by standard breeding practices. All iBIP mice used in this study were on a 70% FVB and 30% mixed background but intercrossed for >20 generations to create a recombinant inbred background suitable for immunocompetent syngeneic tumor establishment. To initiate tumors, iBIP mice were treated with 4-hydroxy-tamoxifen (70% Z-isomer, 30% E-isomer, H6278; Sigma-Aldrich) dissolved in 100% EtOH. One microliter of 100 μM 4-hydroxy-tamoxifen was applied once on the tip of each ear. Mice were then continually administered doxycycline either through the drinking water (2 mg/ml) (D43020; Research Products International) or in the chow (200 mg/kg) (S3888; Bio-Serv). No differences were seen in tumor penetrance or latency between the water and chow vehicles.

For iBIP allografts, iBIP cell lines were established from tumors, cutting the tissues into small pieces (1-3mm in diameter) and digesting them in collagenase IV (100U/ml, #17104019, Thermo Fisher) and DNAse I (10 μg/mL, #07469, StemCell Technologies) at 37C for 2h. Digested cell solutions were filtered through a strainer (100 μm) and cultivated in RPMI, 10%FBS, and 2 mg/ml doxycycline. Cells were detached with Trypsin-EDTA (0.25%) when in the exponential growth curve. Cells were then resuspended in HBSS ($1X10^5$/100μl) and injected intradermally into Nod Scid Gamma (NSG) immunodeficient (Jackson Lab) or syngeneic immunocompetent mice.

"BP" and Yumm1.7 tumors were established as previously described (2-4). Briefly, tumor mouse cells were cultivated in vitro, in RPMI, 10%FBS. Cells were detached with Trypsin-EDTA (0.25%) when in the exponential growth curve. Cells were then resuspended in HBSS ($10^5$/100μl) and injected intradermally in C57BL/6J mice (Jackson Laboratory, Bar Harbor, ME).

For intratumoral injection of chemokines or microparticles, saline was used as a vehicle in a maximum volume of 100μl.

Tumor volumes were calculated using electronic calipers to measure the length (l), width (w), and height (h) and using the formula (l × w × h) × π/6. For all the experiments, when the tumors reached the average volume of 100 $mm^3$, animals were distributed among the treatment groups. PLX4720 chow (417 parts per million, ppm Research Diets Inc.) and its corresponding control diet replaced the standard mouse chow when indicated. MEK162 was resuspended in 1% carboxymethylcellulose/0.5% Tween 80 and administered 7 mg/kg/day by oral gavage using sterile flexible plastic adapters in a volume of 100 µL.

Body mass was measured using an electronic scale. Animals were euthanized when the tumor burden reached >1,000 $mm^3$ or when the tumor became ulcerated. All animal experiments were performed according to protocols approved by the Institutional Animal Care and Use Committees of The University of Texas MD Anderson Cancer Center.

*RNA extraction, expression microarrays, and bioinformatics analysis.* Mouse tumors were collected at the indicated time points and microdissected under a microscope to remove normal skin and cartilage contamination. Tumors were homogenized in TRIzol solution (Thermo Scientific). Nucleic acids were extracted using chloroform, precipitated with isopropanol, washed with 70% ethanol, and resuspended in RNAse-free water. DNA was removed using DNAse in the presence of RNAse inhibitors (Promega) by incubating for 40 min at 37 °C. Samples were processed through the RNEasy kit (Qiagen) to wash and elute the RNA. Samples were tested for purity by measuring 260/230 and 260/280 values on a Nanodrop machine (Thermo Scientific). Impure samples (<1.8 ratio) were repurified using standard ethanol precipitation. Further quality control was performed by the Dana-Farber Cancer Institute Microarray Core facility (http://chip.dfci.harvard.edu/) using the Bioanalyzer platform (Agilent). A minimum of 1 µg of RNA per sample was run on a Mouse Genome 430 2.0 Array (Affymetrix). Microarray data are accessible through GEO (Gene Expression Omnibus) under the accession number GSE79972.

Gene expression data were processed using the affy R package(5), normalized using RMA(6), and differential expression using the limma R package(7).

The Short Time-series Expression Miner (STEM) algorithm(8,9) was used to detect significant trends in genes over time (k-means clustering, k=5). Gene set enrichment using the hypergeometric test and the MSigDB gene set C2.CP.v7.5.1 was performed on all clusters and used to rank gene sets on statistical significance. Gene Set Enrichment Analysis (GSEA) analysis was performed using the pre-ranked program and the MSigDB gene set C5.CP.v7.5.1 gene ontology (G.O.) (10,11). Gene Expression Deconvolution Interactive Tool (GEDIT)(12), using the Minimum Entropy Ranking metric and the Mouse Body Atlas (13).

*TRAP analysis*. The TRAP algorithm has been previously described(14). Here, we have modified the network analysis to focus on cytokines-to-pathway relationships, whereas the original TRAP network focused on transcription factor-to-pathway relationships. Using Biological Process Gene Ontology annotations (GO:0005125, cytokine activity), we first defined a set of 229 putative cytokines. For pathways, we used the canonical pathway gene sets as defined by MSigDB (C2.CP.v7.5.1)(15). With cytokines and pathways defined, we generated the background network model as described in (14). Briefly, using the compendium of mouse gene expression data defined in (14), we calculated pathway expression for each gene set as the average expression of all genes in the gene set (with any of the 229 cytokines removed). Cytokine-to-pathway relationships were calculated using mutual information, and a threshold of 3 standard deviations away from the mean was used to define cytokine-to-pathway relationships. The final TRAP network consisted of 13,629 edges that connected 157 cytokines to 2079 pathways.

The pathways identified from the GO term enrichment using the STEM clustering approach on the immune cluster were used as input to rank cytokine centrality in the TRAP network. A subnetwork of 748 edges was defined by the selected pathways (n=68) and the immediate neighbor cytokines (n=98). A degree centrality measure was used to rank cytokines that had the most edges in the subnetwork compared to the full TRAP network.

*Immunohistochemistry.* FFPE mouse tumor specimens were sectioned with a microtome (5-μm sections) and put on slides. After deparaffinization, citrate-based antigen retrieval was performed. Blocking was performed using Normal Horse Serum (ImmPRESS Reagent Kit, Vector Labs). Slides were then incubated with one of the following primary antibodies overnight at 4°C in a humid chamber: anti Cxcl9 (#701117, clone 11H1L14, Invitrogen), anti Cd8 (#14-0808-82, Clone 4SM15, Invitrogen), anti Cd4 (#14-0041-82, GK1.5, Invitrogen). Slides were then incubated with HRP-conjugated anti-rabbit secondary antibody. Processed slides were finally added with TSA Plus Fluorescein System (Perkin-Elmer) to allow fluorophore deposition and counterstained with hematoxylin (Vector Laboratories) and DAPI. Coverslips were mounted after dehydration of the sections using Permanent Mounting Medium (Vector Laboratories). Images were digitally acquired with a Nikon Eclipse T2 microscope connected to DS-Ri2 camera (Nikon). Quantification of IHC expressed as the number of positive cells/ number of total cells was performed using ImageJ (Rasband, W.S., ImageJ, U.S. National Institutes of Health, Bethesda, MD, https:// imagej.nih.gov/ij/, 1997–2016).

*Immunofluorescence.* Immunohistochemistry. FFPE mouse tumor specimens were sectioned with a microtome (5-μm sections) and put on slides. After deparaffinization, citrate-based antigen retrieval was performed. Blocking was performed using Normal Horse Serum (ImmPRESS Reagent Kit, Vector Labs). Slides were then incubated with the following primary antibodies overnight at 4°C in a humid chamber: anti Cxcr3 (#LS-B10183, LSBio), anti Cd8 (#14-0808-82, Clone 4SM15, Invitrogen). Slides were then incubated with fluorescence-conjugated anti-rabbit or anti-rat secondary antibodies (Alexa Fluor 488, Alexa Fluor 647, ThermoScientific). Processed slides counterstained with hematoxylin (Vector Laboratories). Slides were processed with

TrueVIEW Autofluorescence Quenching Kit, and coverslips were mounted using VECTASHIELD Vibrance Antifade Mounting Medium with DAPI. Images were digitally acquired with the ImageXpress Pico Imaging system (Molecular Devices). Quantification was performed using the CellReporterXpress Image Acquisition and Analysis Software (Molecular Devices).

*Patients' RNAseq data.* Melanoma patients' RNAseq methods and data were previously published(1). Patient samples were exhausted during the analysis.

*Chemotaxis and activation assays.* EasySep™ Mouse Cd8+ T Cell Isolation Kit (StemCell Technologies) was used according to the manufacturer's instructions to derive Cd8 T-cells from the spleens of the animals. Isolated T-cells were then cultured in plates pre-coated with anti-Cd3 (0.5 µg/ml, #100339, Clone 145-2C11, Biolegend) and anti-Cd28 (5 µg/ml, #102115, clone 37.51, Biolegend), for lymphocyte activation. The Cd8 + T cells were cultured in RPMI1640 medium containing 10% heat-inactivated FBS and supplemented with L-glutamine (2 mM), penicillin (50 U/ml), streptomycin (50 µg/ml), 2-mercaptoethanol (50 µM), and IL-2 (30 U/ml, #212-12, PeproTech).

Following 72h of activation, Cd8 lymphocytes were assayed through a chemotaxis experiment in a Boyden chamber (5µm pores, 24 well plates). Briefly, lymphocytes were counted and distributed in the top chambers. Then, recombinant chemokines or microparticle-soaked medium were added to the bottom chamber at the desired concentration. Medium and FBS concentration was the same for the bottom and top chamber (RPMI1640 medium + 10% FBS). After 3h, bottom and top chamber media were collected, and the number of cells was counted through a Guava Flow Cytometer. The migration index was calculated as the number of cells migrated to the bottom chamber/total number of cells and normalized to the control group.

For T-cell activation assessment, T Cd8+ cells were incubated in the presence of the indicated recombinant mouse chemokines (rCxcl9 #250-18, rCxcl10 #250-16, rCxcl11 #250-29, PeproTech) for 72h, collected and analyzed through flow cytometry.

*Porous Silica Nanoparticle Synthesis.* Cetyltrimethylammonium bromide (CTAB, 141.75 mg) was dissolved in 70ml d$H_2O$ in a 250 ml screw-cap container. Ammonium hydroxide ($NH_3 \cdot H_2O$, 3.03 ml) (28-30%) was added dropwise into the reaction flask. The mixture was stirred for 1h at 1200 rpm. Tetraethyl orthosilicate (TEOS, 0.6025 ml) was added to the mixture and stirred vigorously for 4h. The mixture was centrifuged at 5000 rpm for 10 minutes. The particles were washed and dispersed in EtOH and $H_2O$ 3 times (ratio 1:1 v/v). The mixture was centrifuged at 5000 rpm for 10 minutes between each wash. The final solution was biphasic. The particles were dispersed in 20 ml of 1:1 1N acetic acid: dichloromethane 3 times. The mixture was washed in $H_2O$ 2 times and centrifuged at 5000 rpm for 10 minutes between each wash. The mixture was resuspended in Isopropyl alcohol (IPA) for storage. When ready to be processed, the mixture was then placed in a vacuum oven at 60°C and -36 Pa to dry before proceeding with loading and encapsulation with PLGA 50:50 (see below). Size and Polydispersity Index (PDI) were measured using Dynamic Light Scattering (DLS). The physical characteristics of silica particles were evaluated using Transmission electron microscopes (TEM) and scanning electron microscopes (SEM) (**Fig. S4 B-C**), revealing the characteristic porous structure. Dynamic light scattering (DLS) reported an average size of 371±5.7 nm and a Polydispersity index (PDI) of 0.093 ±0.027 (**Fig. S4D-E**), indicating that nanoparticles were homogenous and uniform. We then included the silica core in a PLGA 5% or 10% shell (**Fig. S4F-G**), which increased the average particle size to 1003 nm±47 (**Fig. S4H**). The average rCxcl9 encapsulation efficiency was 39.5±0.9 % (**Fig. S4I**).

*Loading of Silica Nanoparticles with chemokines.* The desired amount of dried silica nanoparticles was placed into a 1.5ml Eppendorf tube. The loading solution was prepared with 2.5 μg (in vitro experiments) or 50µg (in vivo experiment) of rCxcl9 (with BSA carrier, #250-18, Preprotech) in 500 µl PBS. The particles in solution were placed on an orbital mixer at 37°C for 2 hrs. Particles were centrifuged to separate them from the solution (5000 rpm for 10 min). The loaded particles were dried in a lyophilizer for 3-8h.

*Porous Silica Nanoparticle encapsulation with PLGA 50:50.* A water solution (2.5% and 1% PVA) was prepared as follows: (1) 5% PVA stock solution was prepared by placing 25g PVA (13.000-23.000 MW), and 500ml DI water in a screw-top Glass Container and autoclaved. (2) 5% stock PVA was diluted to 2.5% PVA and 1% PVA with DI water. An oil solution (10% or 5%wt PLGA 50:50/DCM) was prepared as follows: (1) 100mg (105 solution) or 50mg (5% solution) of PLGA were placed (50:50, viscosity range 0.76-0.94 dL/G, lot#A17-065, Durect Corporation) in a scintillation vial with 2ml DCM.

Emulsion #1 was then performed by placing the oil solution into the homogenizer tube. Silica particles were added to the oil Solution. In the homogenizer tube, 3ml of 2.5% PVA were added, and the solution was homogenized for 5 mins at 1200 rpm. Emulsion #2 was then performed by placing Emulsion #1 (5ml) in Water solution (40ml, 1% PVA) in a 100ml beaker on a stir plate, covering it, and stirring at 600 rpm for a minimum of 6h (or overnight). Contents of the beaker were transferred into 50ml falcon tubes and centrifuged at 5000 rpm for 10 mins. The supernatant was removed. Particles were then transferred to clean 1.5 ml lock-cap Eppendorf tubes, frozen for 1h at -80°C, then lyophilized for a minimum of 4h. Particles were stored at -80°C until ready to be used. Imaging of particles was recorded using Scanning Electron Microscope (SEM) and Transmission electron microscopes (TEM). The specific PLGA-silica formulation was selected as the PLGA copolymer is approved by the Food and Drug Administration and European Medicines Agency for drug delivery applications.

*Flow Cytometry.* Mouse tumors were collected, cut into small pieces (1-3mm in diameter), and digested with a mixture of Collagenase IV (100U/ml, #17104019, Thermo Fisher) and DNAse I (10 μg/mL, #07469, StemCell Technologies). Tumors were incubated for 2h at 37°C, centrifuged, and filtered through a 100μm strainer. Red Blood Lysis solution was applied (#118-156-101, Quality Biological). Cells were then resuspended at a density of $1X10^6$/ml in flow cytometry binding buffer and stained with the following fluorophore-conjugated antibodies (Biolegend): Cd8 (clone 53-6.7), Cxcr3 (clone CXCR3-173), Cd69 (clone H1.2F3), Cd25 (clone PC61). 7-AAD Viability Staining Solution was used following the manufacturer's instructions (Biolegend). Guava easyCyte 11HT Benchtop Flow Cytometer was utilized for the acquisition, and results were analyzed through FlowJo™ v10.8 Software (BD Life Sciences). The gating strategy for mouse samples flow-cytometric analysis is reported in **Fig. S10**.

*Graphic elements.* The schematics in the paper were created with BioRender.com (agreement number MU24MQR7RN). The statistical graphs were created with Graphpad Prism v9 (GraphPad Software LLC).

### SUPPLEMENTARY FIGURE LEGENDS

**Figure S1. Minimal Residual Disease contains residual tumor cells.** Immunohistochemical staining for GFP, a built-in tumor marker in the R26-lsl-rtTA-IRES-GFP allele. DAPI is used as a nuclear counterstain. Bars represent 100 μm.

**Figure S2. Cxcr3 ligands differentially induce early and late T-activation markers.** Flow cytometric assessment of early Cd69 A) and late Cd25 B) activation markers induced by the indicated recombinant chemokines (100 ng/ml) after 72h of treatment. Data are indicated as average ± SEM and expressed as the percentage of positive cells over the total Cd8+ T cells.

**Fig.S3. Cxcl9 is induced in consequence of pharmacologic BRAFi in mouse tumors.** Cd8 isoforms Cd8b1 and Cd8a and Cxcl9 RNA expression of "BP" tumor sections in consequence of pharmacologic BRAFi (PLX4720, 417 parts per million, ppm, 1 week of treatment, RNAseq; n=3).

**Fig S4. Recombinant Cxcl9 delays the occurrence of tumor relapse in iBIP allografts.** Iindividual tumor growth curves of iBIP allografts injected with rCxcl9 or vehicle (Ctrl, n=5, rCxcl9, n=7) after dox withdrawal. Indicated tumor volumes are relative to the volume at MRD establishment (24d), when tumors were rechallenged with dox and observed for relapse.

**Fig.S5 Characterization of Silica-PLGA microparticles.** A) Graphical representation of Silica-PLGA nanoparticles fabrication process. B) High magnification TEM image of a silica nanoparticle. Scale bar 50 nm. C) SEM image of silica nanoparticles. Scale bar 1μm. D) Silica nanoparticle average size as determined by DLS (Dynamic light scattering). E) Silica nanoparticle Polydispersity index (PDI) as determined by DLS. F) High magnification SEM image of silica nanoparticles encapsulated in PLGA 10% shell. Scale bar 5μm. G) Low magnification SEM image of silica nanoparticles encapsulated in PLGA 10% shell. Scale bar 20μm. (I) Silica-PLGA loaded with Cxcl9 encapsulation efficiency. Data are represented as mean ± SD (n=3).

**Fig. S6. Microparticle-delivered recombinant Cxcl9 prolongs BRAFi efficacy in Yumm1.7 allografts.** Individual tumor growth curves of Yumm1.7 allografts treated with PLX4720 with or without microparticles that contained BSA or rCxcl9 (n=10). Indicated tumor volumes are relative to the volume at MRD establishment (24d).

**Fig. S7. BRAFi+MEKi+rCxcl9 causes complete pathologic response in Yumm1.7 tumors.** Representative Hematoxylin and Eosin staining of Yumm1.7 tumors treated PLX4720 or PLX4720+MEK162 with or without microparticles that contained BSA (mBSA) or rCxcl9 (mrCxcl9), n=5. T=tumor; NT=non-tumor. Images were all acquired with a 10X objective. Bars represent 150 μm.

**Fig. S8. Microparticle-delivered rCxcl9 in combination with BRAFi induces Cxcr3+ Cd8+ T-cell infiltration.** Representative images of double immunofluorescence staining for Cd8 and

Cxcr3 (n=5 tumors, whole section acquired, 10x objective). DAPI (blue) is used as a nuclear counter stain. Cd8+ cells are in red (Alexa Fluor 647), Cxcr3+ cells are in green (Alexa Fluor 488). Bars represent 70 µm.

**Fig. S9. Microparticle-delivered rCxcl9 in combination with BRAFi induces Cd8+ T-cell infiltration.** Immunofluorescence quantification of Cd8+ T cell abundance in Yumm1.7 tumors treated with PLX4720 or PLX4720+MEK162 with or without microparticles that contained BSA or rCxcl9 (n=5). The percentage of Cd8+ T cells is calculated over the total number of cells in the tissues.

**Fig S10. Gating strategy for flow cytometric analysis.** The gating strategy for the detection of CD8+CXCR3+ in mouse samples is reported. After debris exclusion (1), doublets elimination (2), and viability assessment (3), the percentage of negative, single positive, and double positive cells were assessed (4).

### SUPPLEMENTARY TABLE LEGENDS

**Table S1. k-means clustering results of gene expression data.** The average gene expression for each time point was calculated, then the k-means clustering algorithm (k=5) was applied using the STEM software.

**Table S2. Gene set enrichment results for the immune gene expression cluster.** Enrichment analysis was performed for the gene expression cluster that was predominantly immune-related genes. The hypergeometric test was applied using the genes in the cluster and the C2.CP.v7.5.1 collection of gene sets from the MSigDB.

**Table S3. Cytokines ranking based on their gain in centrality from the TRAP analysis for the immune gene expression cluster.** Gene sets that were statistically significantly enriched in the immune gene expression cluster were selected from the TRAP network. These gene sets and the immediate neighbors, which are cytokines, define the immune gene expression cluster subnetwork. Cytokines were then ranked based on their centrality in the subnetworks compared to the full TRAP network.
