## Supplementary figures and images for "Microparticle-delivered Cxcl9 prolongs Braf inhibitor efficacy in melanoma"

### FigS1-5

**Fig. S1**

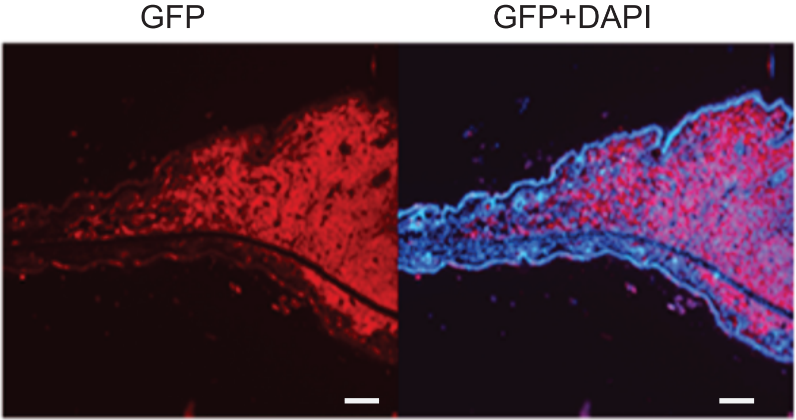

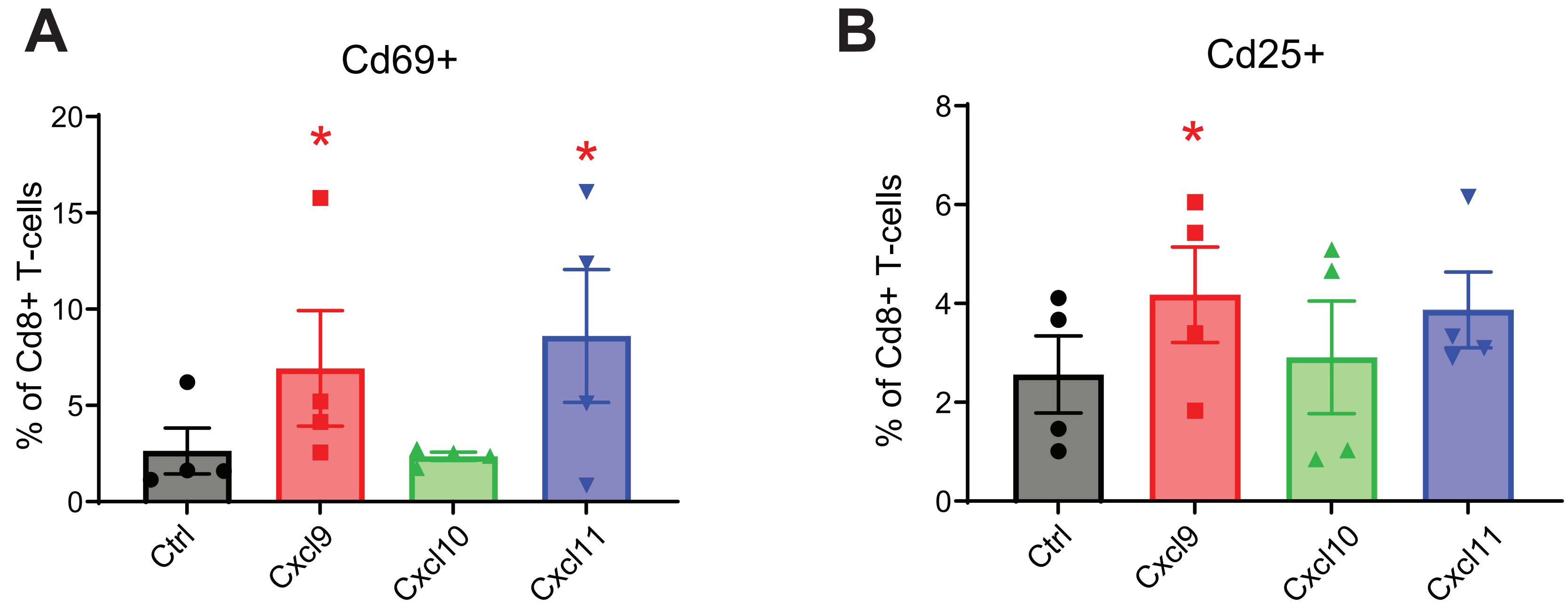

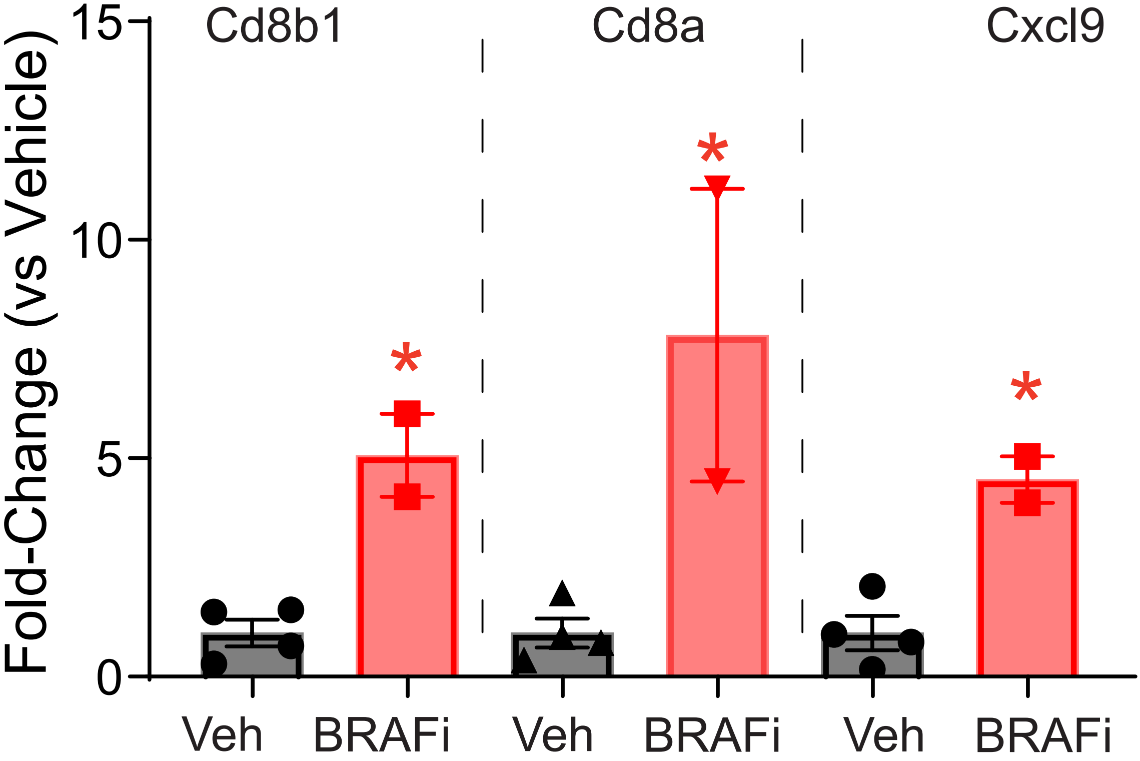

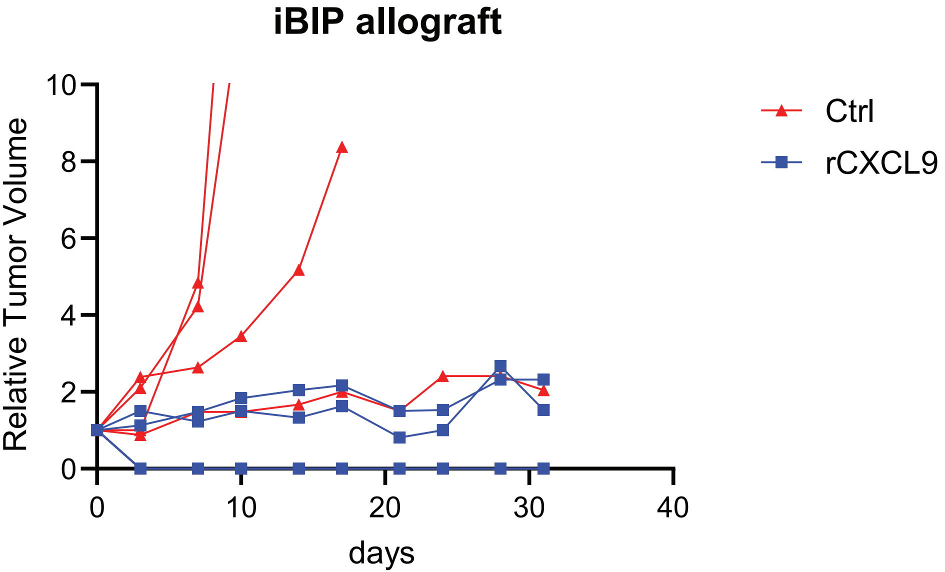

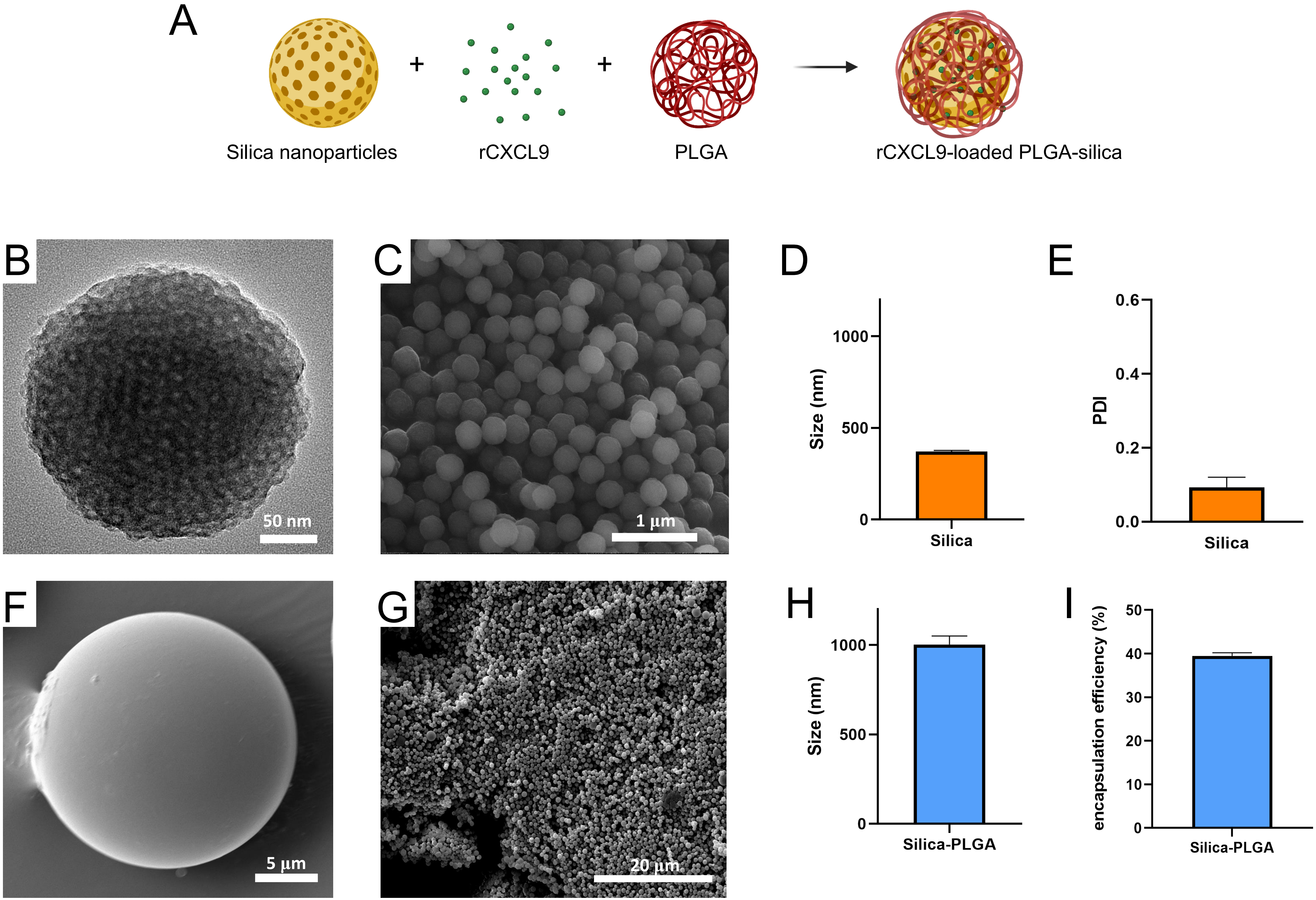

### FigS6-10

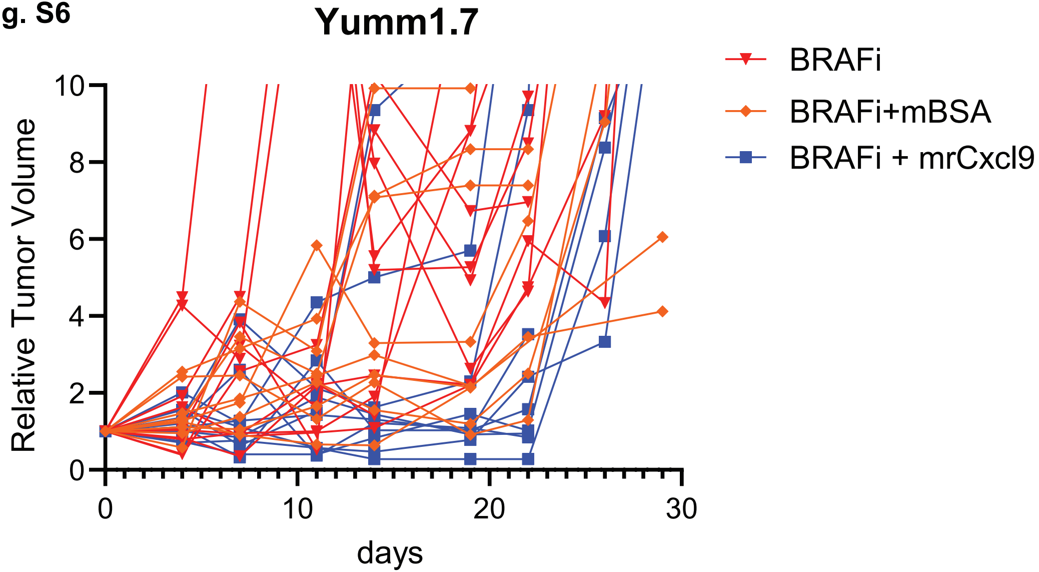

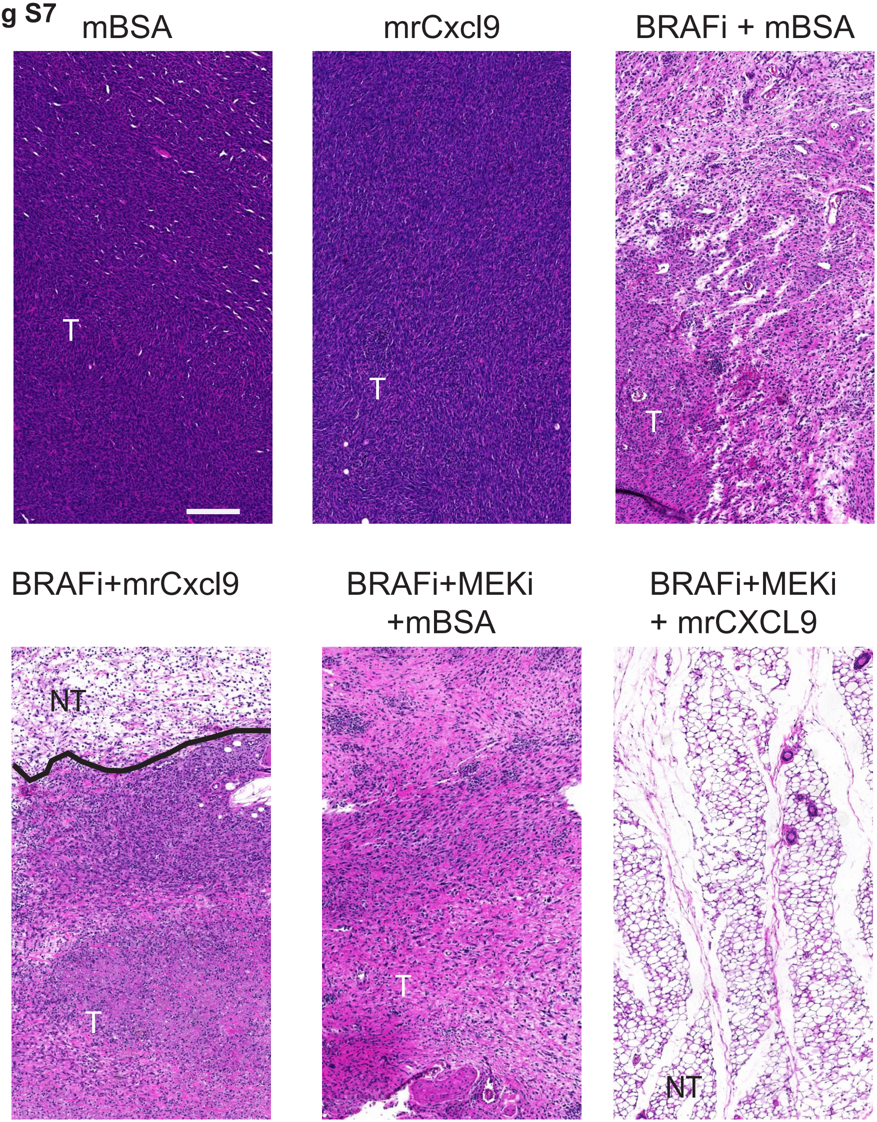

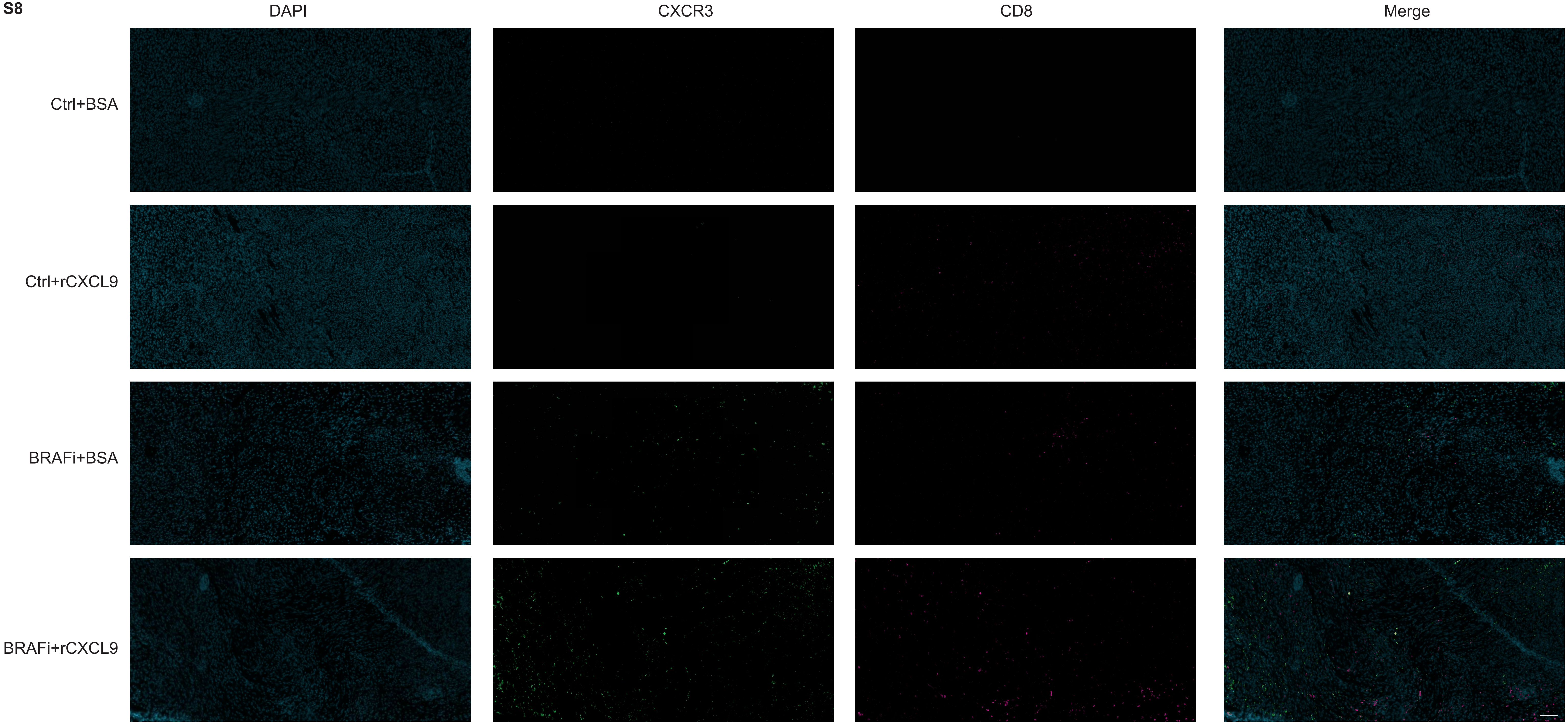

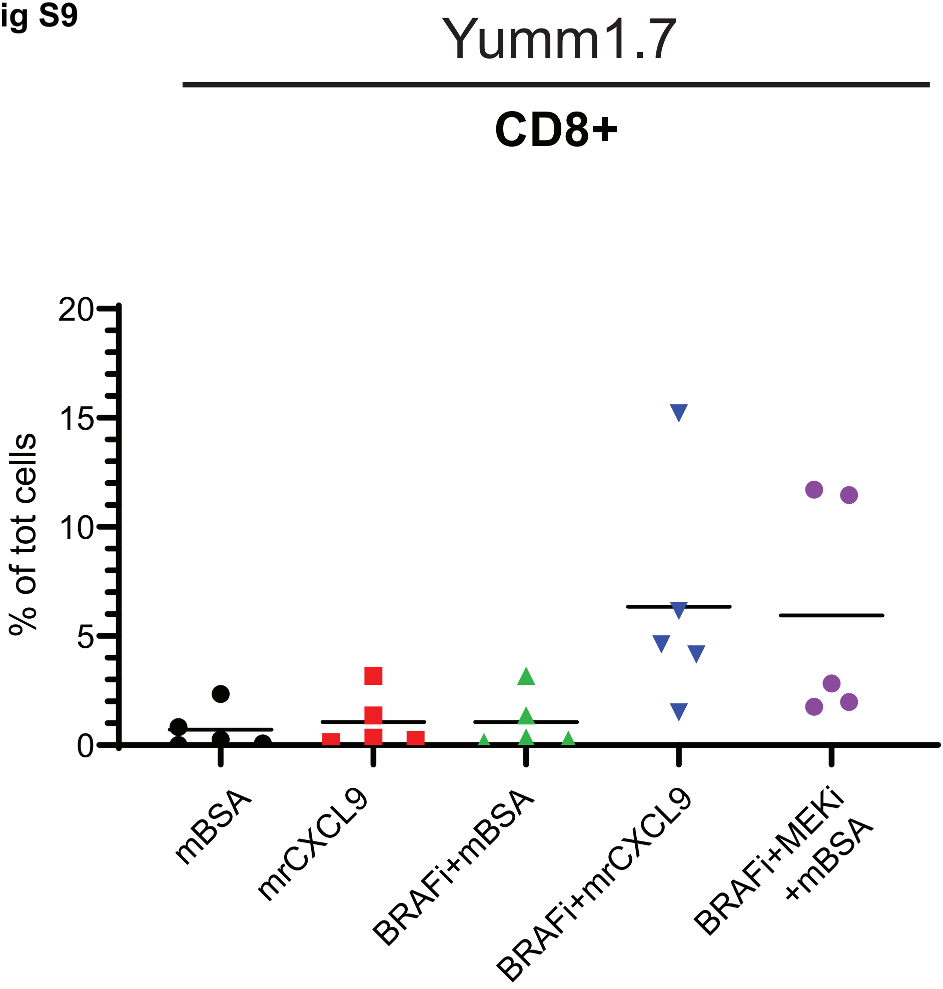

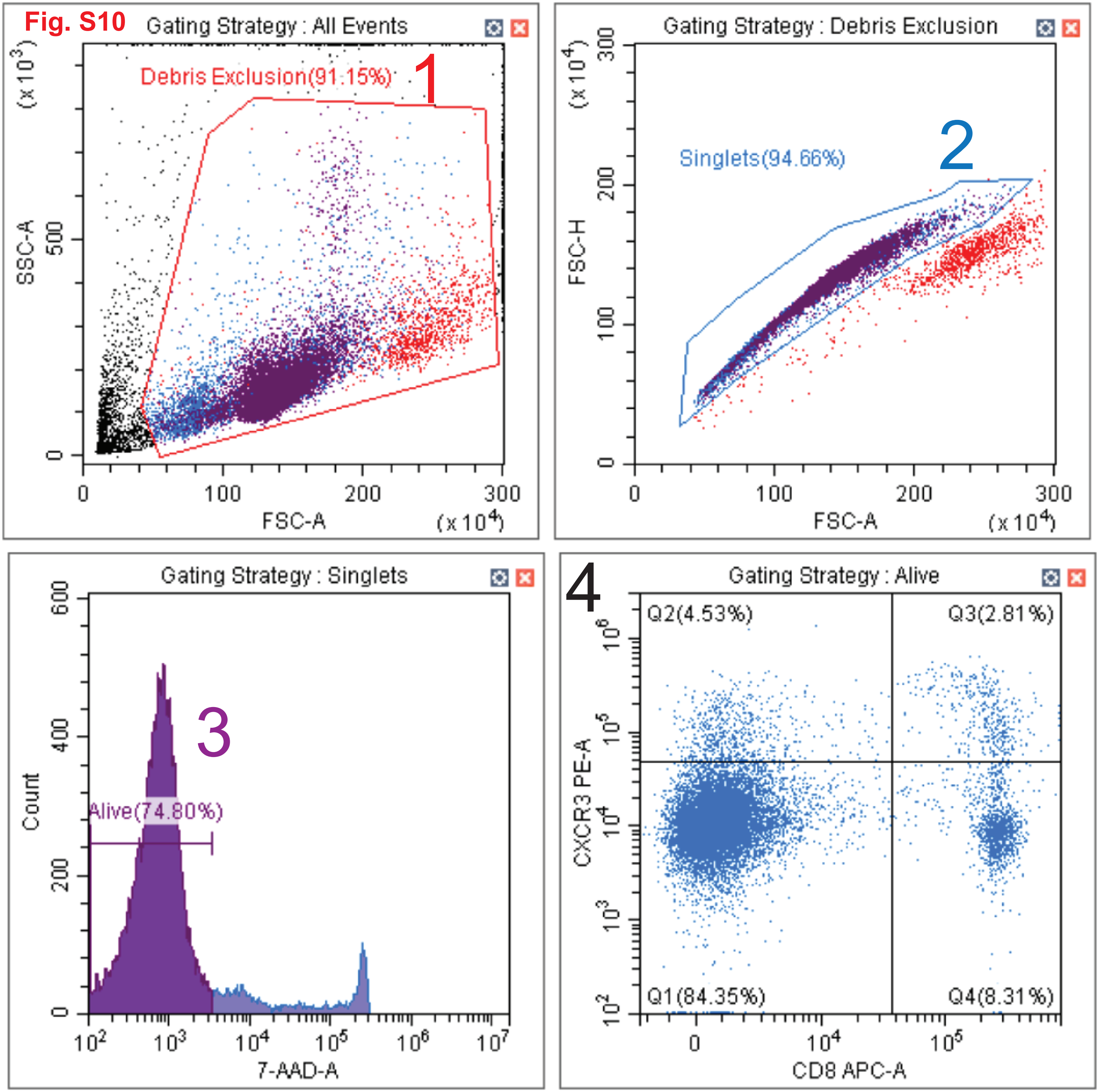
